# Transcriptional repression of Myc underlies AGO1’s tumour suppressor function

**DOI:** 10.1101/2020.03.10.984906

**Authors:** Olga Zaytseva, Naomi C. Mitchell, Linna Guo, Owen J. Marshall, Linda M. Parsons, Ross D. Hannan, David L. Levens, Leonie M. Quinn

## Abstract

Here we report novel tumour suppressor activity for the *Drosophila* Argonaute family RNA binding protein AGO1, a component of the miRNA-dependent RNA-induced silencing complex (RISC). The mechanism for growth inhibition does not, however, involve canonical roles as part of the RISC; rather AGO1 controls cell and tissue growth by functioning as a direct transcriptional repressor of the master regulator of growth, Myc. AGO1 depletion in wing imaginal discs drives a significant increase in ribosome biogenesis, nucleolar expansion, and cell growth in a manner dependent on Myc abundance. Moreover, increased *Myc* promoter activity and elevated *Myc* mRNA in AGO1 depleted animals requires RNA Pol II transcription. Further support for transcriptional AGO1 functions is provided by physical interaction with the RNA Pol II transcriptional machinery (chromatin remodelling factors and Mediator Complex), punctate nuclear localisation in euchromatic regions and overlap with Polycomb Group transcriptional silencing loci. Moreover, significant AGO1 enrichment is observed on the *Myc* promoter and AGO1 interacts with the *Myc* transcriptional activator Psi. Together our data show AGO1 functions outside of the RISC to repress *Myc* transcription and inhibit developmental cell and tissue growth.

## Introduction

Tightly coordinated regulation of cell and tissue growth is essential for animal development; decreased growth leads to small organs and diminished body size, while heightened proliferative growth is associated with genomic instability and cancer. The MYC transcription factor and growth regulator has been studied extensively since identification as an oncogene in the early eighties (Vennstrom et al., 1982), and given MYC overexpression due to chromosomal translocation directly drives malignant transformation in Burkitt’s Lymphoma (Dalla-Favera et al., 1982; Taub et al., 1982). Research in subsequent decades implicated increased MYC in progression of most tumours (Dang, 2012; Liao and Dickson, 2000; Meyer and Penn, 2008). In normal adult tissues, MYC expression is relatively low and generally restricted to cells with regenerative and proliferative potential (Marcu et al., 1992). Even small increases in MYC abundance are sufficient to promote proliferative cell growth (reviewed in (Dang, 2010; Levens, 2010; Zaytseva and Quinn, 2017)), thus, understanding the molecular control of *MYC* expression will provide critical insight into mechanisms of MYC dysregulation in cancer.

In normal cells, MYC is regulated by signalling inputs from a diverse array of developmental and growth signalling pathways (Zaytseva and Quinn, 2017). The many cellular signalling inputs converging on *MYC* transcription are integrated by FUBP1, a KH domain protein that binds single stranded DNA and interacts with the general transcription factor complex TFIIH to modulate *MYC* promoter output (Chung and Levens, 2005; Chung et al., 2006; He et al., 2000; Liu et al., 2006; Zhang and Chen, 2013). The mammalian FUBP family comprises 3 proteins (FUBP1-3) (Zhang and Chen, 2013), which are represented by one ortholog in *Drosophila*, Psi. Like FUBP1, Psi also interacts with RNA Pol II transcriptional machinery, particularly the transcriptional Mediator (MED) complex, to pattern *Myc* transcription, cell and tissue growth in the *Drosophila* wing epithelium (Guo et al., 2016). In addition to roles in transcription, Psi binds RNA via the KH domains and interacts with the spliceosome to regulate mRNA splicing (Labourier et al., 2001; Wang et al., 2016). Although Co-IP Mass Spectrometry detected Psi in complex with the Argonaute protein, AGO1 (Guo et al., 2016), the potential significance of this interaction is unknown.

Argonaute proteins comprise the core of the RNA-Induced Silencing Complex (RISC), which uses noncoding RNA as a guide to target mRNAs for post-transcriptional gene silencing. *Drosophila* AGO2 is best characterised as part of the siRNA-induced silencing complex (siRISC) (Okamura et al., 2004), while AGO1 predominantly functions in microRNA-induced silencing complexes (miRISCs) and post-transcriptional mRNA silencing (Förstemann et al., 2007). Of importance to this study, AGO1-mediated mRNA silencing has been implicated in transcript destabilisation and translational repression of *Myc* in flies (Daneshvar et al., 2013) and humans (Challagundla et al., 2011). Here, however, we report a novel role for AGO1 as a direct *Myc* transcriptional repressor and demonstrate that this underlies cell growth inhibition. AGO1 depletion not only increases *Myc* promoter activity, mRNA and protein abundance, but the elevated *Myc* expression requires RNA Pol II transcriptional activity. Localisation to the nucleus, together with interaction with transcriptional machinery and significant AGO1 enrichment on the *Myc* promoter suggests, in addition to the established roles in miRNA silencing in the cytoplasm, AGO1 constrains *Myc* transcription to control cell and tissue growth during development.

## Results

### AGO1 interacts with Psi and RNA Pol II transcriptional machinery

The single stranded DNA/RNA binding protein Psi has essential roles in *Myc* transcriptional control and RNA processing in *Drosophila*. In addition to physically and genetically interacting with the RNA Pol II transcriptional machinery, the *Drosophila* Protein Interaction Map (DPiM) large scale co-IP mass-spectrometry (Guruharsha et al., 2011) suggested association between Psi and AGO1 (Guo et al., 2016). Our analysis of the DPiM identified Psi as the most frequent AGO1 interacting partner (**Figure 1A**). Ontological class analysis for the top 70 AGO1 interactors revealed RNA processing factors (49%), as expected, however most (59%) interactors had ascribed functions in RNA Pol II transcription (**Figure 1A-C**, note 10 factors are implicated in both transcription and RNA processing). As the DPiM studies were performed *in vitro* with overexpressed tagged proteins in *Drosophila* S2 cell lines, we validated the interaction between endogenous AGO1 and Psi *in vivo* using Co-IP from wild type 3rd instar larval imaginal tissue. Immunoprecipitation using anti-Psi antibody, followed by anti-AGO1 Western detected a 110 kDa band for AGO1 (**Figure 1D**), while reciprocal IP with anti-AGO1 antibody precipitated the 97 kDa Psi band (**Figure 1E**). The observation that endogenous AGO1 and Psi form a complex *in vivo* led us to investigate potential genetic interactions between AGO1 and Psi.

**Figure 1.**
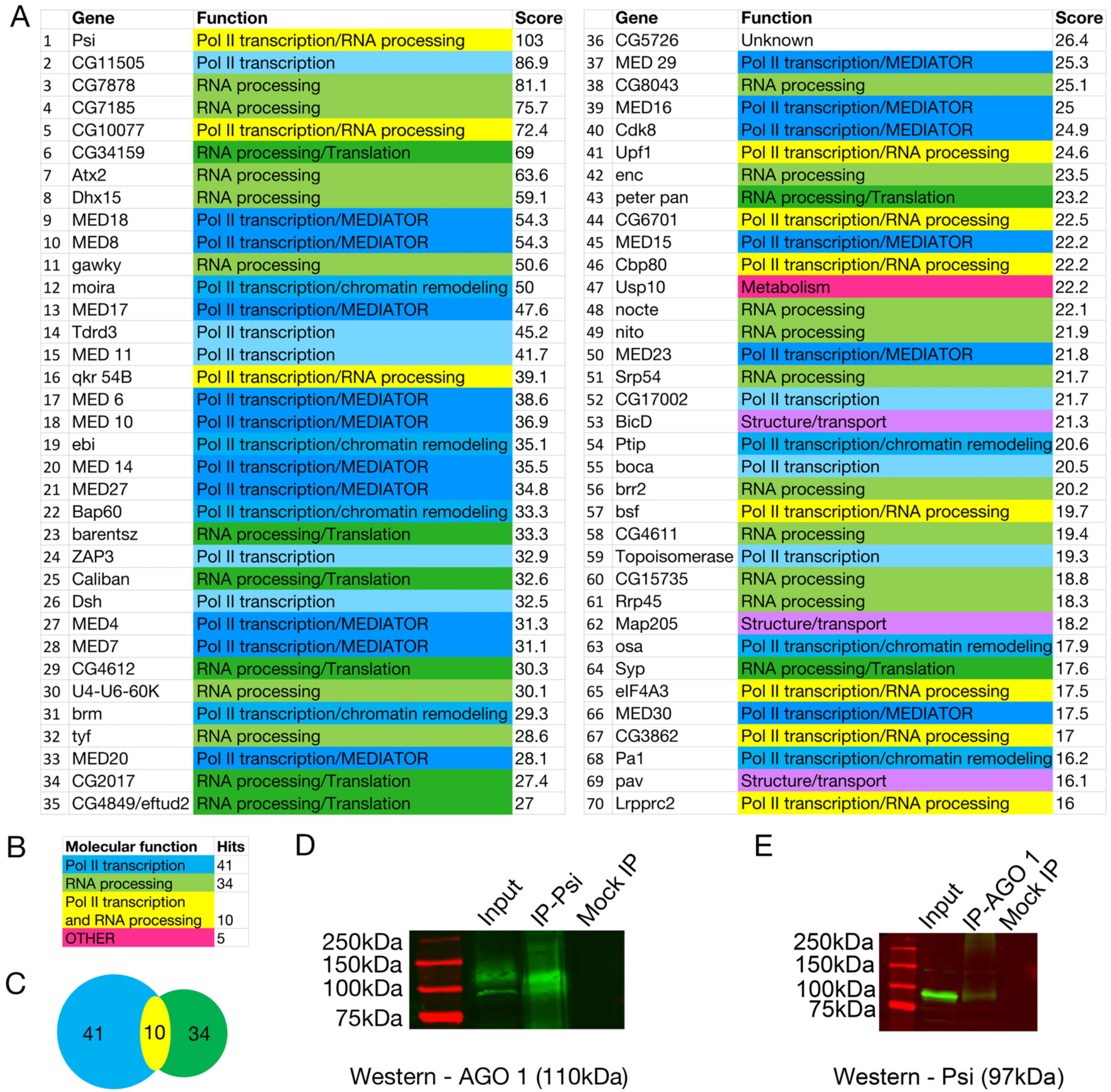
AGO1 interacts with RNA Pol II transcriptional machinery and RNA processing factors. **(A)** List of top 70 AGO1 interactors from the *Drosophila* Protein Interaction Map (DPiM) data set. **(B)** Summary of ontology classes for the top 70 AGO1-interactors. **(C)** Intersection of interactors with functions in RNA Pol II transcription and/or RNA processing. **(D, E)** Co-IP of endogenous Psi and AGO1 from wild type third instar larvae. **(D)** IP with anti-Psi and Western blot with anti-AGO1 (110 kDa). **(E)** IP with anti-AGO1 and Western blot with anti-Psi (97 kDa).

### Psi-dependent growth is sensitive to AGO1 abundance

Psi knockdown in the dorsal wing compartment results in a “wings up” phenotype as impaired cell and tissue growth of the top layer of the wing results in torsional strain and wing bending (Guo et al., 2016). We therefore tested whether this was modulated by AGO1 depletion, using two independent P-element insertion mutants (*AGO1*^*k00208*^ and *AGO1*^*04845*^). Interestingly, *AGO1* heterozygosity alone was sufficient to increase wing size, suggesting AGO1 normally constrains growth. Moreover, heterozygosity for either *AGO1* mutant suppressed impaired tissue growth caused by Psi depletion (**Figure 2A-C**). Thus, AGO1 normally functions as a negative growth regulator and the wing overgrowth associated with AGO1 reduction is dependent upon Psi.

**Figure 2.**
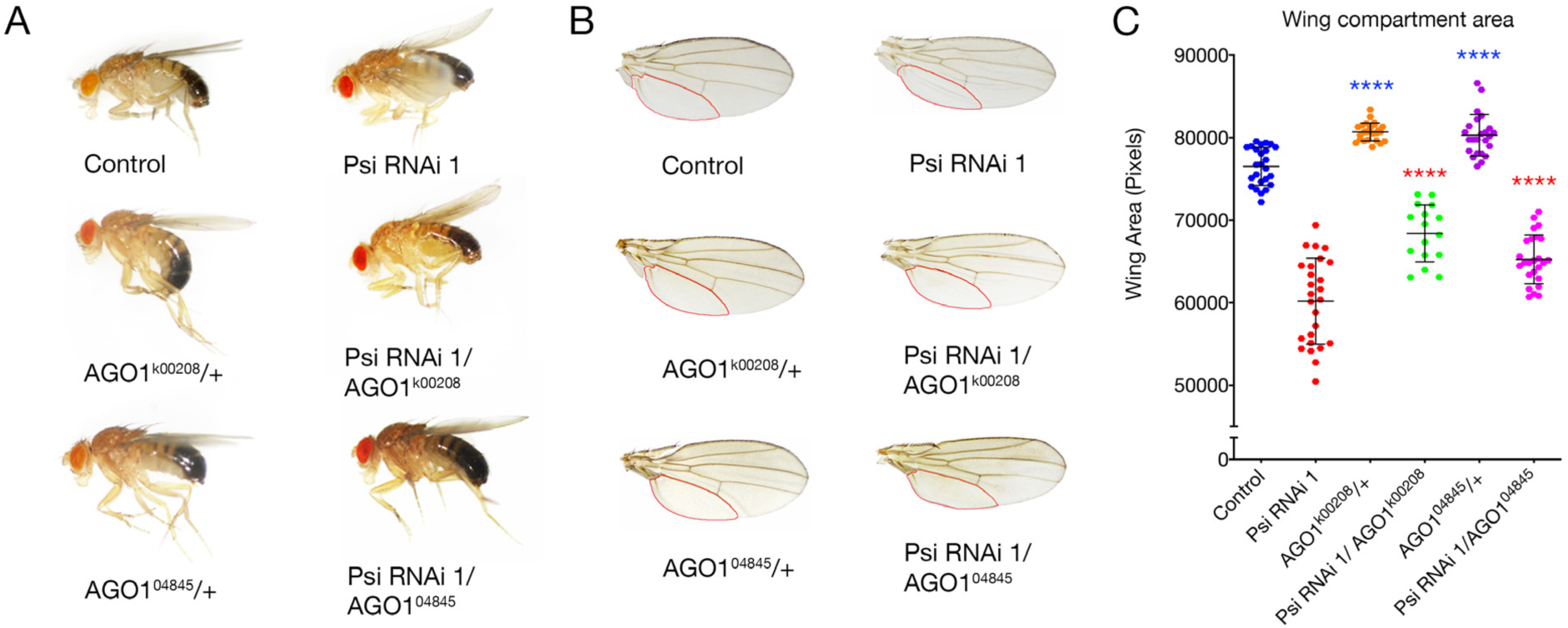
Impaired cell and tissue growth in the Psi knockdown wing is sensitive to AGO1 levels. **(A)** Adult flies and **(B)** wings, compartment below vein L5 outlined in red (genotypes as marked) **(C)** Quantification of compartment area below vein L5 in adult wings, p<0.0001 for the *AGO1* mutant compared with control and the *AGO1* mutant in the *ser-GAL4*>*Psi* RNAi background compared with Psi knockdown alone.

### AGO1 depletion drives cell growth in a Myc-dependent manner

To further examine the cellular basis of the tissue overgrowth associated with AGO1 depletion in specific compartments of the larval wing, we used two independent non-overlapping *AGO1* RNAi lines. We first demonstrated efficient mRNA knockdown for both *AGO1* lines in the wing (**Supplementary Figure 1A**) and dorsal compartment-specific protein knockdown 24 hours after induction of *ser-*GAL4 (**Supplementary Figure 1B**). As pupal lethality and dorsal compartment cell death were associated with constitutive *ser-*GAL4 driven AGO1 knockdown (**Supplementary Figure 2**) the baculoviral caspase inhibitor p35 was co-expressed to prevent apoptosis (Hay et al., 1994) and enable investigation of potential changes to cell growth.

Cell growth requires ribosomal RNA (rRNA) synthesis, processing and assembly with ribosomal proteins (RPs) into 40S and 60S ribosomal subunits in the nucleolus. Thus, the size of this structure, measured by nucleolar-specific fibrillarin antibody, provides an indirect measure of ribosome biogenesis (Mitchell et al., 2015). AGO1 depletion significantly increased nucleolar size (**Figure 3A, B**), suggesting AGO1 normally functions to inhibit cell growth. Consistent with AGO1 depletion driving nucleolar expansion, at least in part due to increased rDNA transcription, transient *AGO1* knockdown significantly increased pre-ribosomal RNA (rRNA) levels and *Polr1c* (RNA Polymerase 1 subunit) mRNA (**Figure 3C**). AGO1 depletion also significantly increased levels of the ribosomal protein subunits *RpS19* and *RpS24* (**Figure 3C**). Together these data suggest AGO1 normally inhibits ribosome biogenesis and cell growth in the wing imaginal disc. ChIP-sequencing studies have previously identified AGO2 binding throughout the 47S region of the human rRNA gene in human cell lines (Atwood et al., 2016), suggesting direct roles for AGO proteins in rDNA transcription and/or rRNA processing. However, these observations would not explain the increase in expression of RNA Pol II-transcribed ribosomal proteins and RNA Pol I subunits, nor the increase in overall cell growth, which requires coordinated activity from all 3 RNA polymerases: RNA Pol I, II and III (transcription of 5S rRNA).

**Figure 3.**
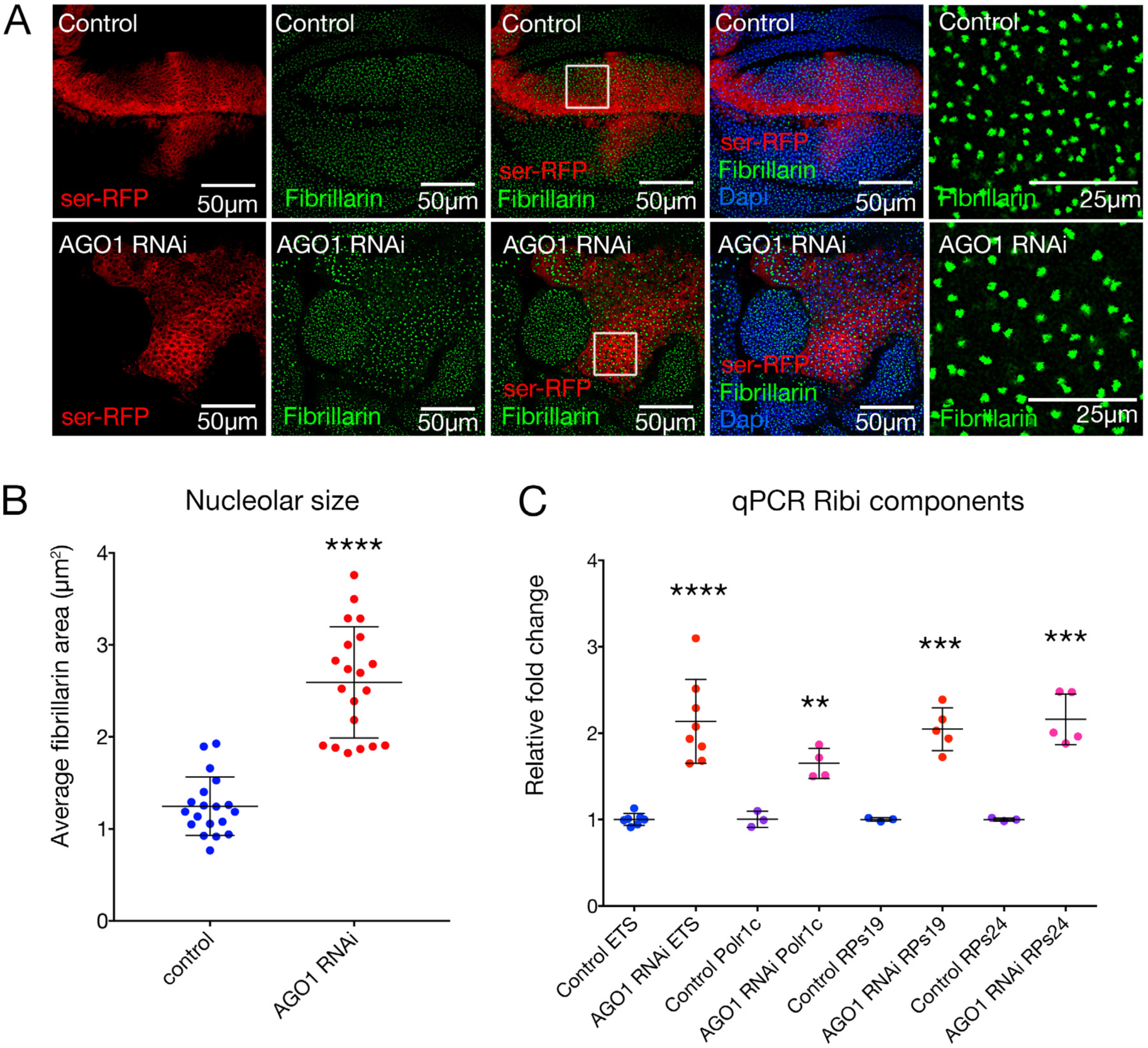
AGO1 knockdown increases ribosome biogenesis. **(A)** Third instar wing discs with *ser*-GAL4 driven *AGO1* RNAi in the *UAS-p35* background compared with the *p35* alone control, marked with *UAS-RFP* and stained with anti-fibrillarin (green) and DNA (blue). Zoom of region in the white square in the panel on the far right. **(B)** Quantification of average nucleolar area of approximately 50-100 nucleoli within the dorsal compartment, taken from a confocal z-section through the wing, p<0.0001 for *AGO1* RNAi 1 compared with control. **(C)** qPCR for 47S pre-RNA and ribosomal proteins (*RpS19* and *RpS24*) and RNA Pol I subunit (*Polr1c*). AGO1 knockdown in the wing significantly increased abundance of *Polr1c* (p<0.01), 47S rRNA 5’ ETS (p<0.0001), *RpS19* and *RpS24* (p=0.0004 and p=0.0006, respectively).

MYC drives cell growth by stimulating transcription of all three RNA Polymerases to upregulate ribosome production (Poortinga et al., 2014). MYC directly stimulates the initiation of RNA Pol I-mediated transcription in mammals (Arabi et al., 2005; Grandori et al., 2005; Shiue et al., 2009), activates transcription of RNA Pol II-transcribed genes encoding the ribosomal proteins (RPs), rRNA processing factors, and components of the nucleolus essential for ribosome biogenesis (Grandori et al., 2005; Grewal et al., 2005; Poortinga et al., 2011). Furthermore, MYC directly activates RNA Pol III transcription to increase 5S rRNA expression, for assembly of the large 60S ribosomal subunit, and tRNA for translation of mRNA into protein (Fernandez, P. C. et al., 2003; Gomez-Roman et al., 2006; Oskarsson and Trumpp, 2005). Myc depletion reduced nucleolar expansion in AGO1 knockdown wing cells down to the control range and co-depletion of the Myc-regulator Psi also significantly decreased nucleolar size (**Figure 4A-B**). Thus, the increased ribosome biogenesis and cell overgrowth associated with AGO1 depletion is dependent on the Psi-Myc axis. The observation that AGO1 depletion increases ribosome biogenesis in a Myc and Psi-dependent manner was consistent with heterozygosity for *AGO1* increasing growth in the Psi knockdown background (**Figure 2**).

**Figure 4.**
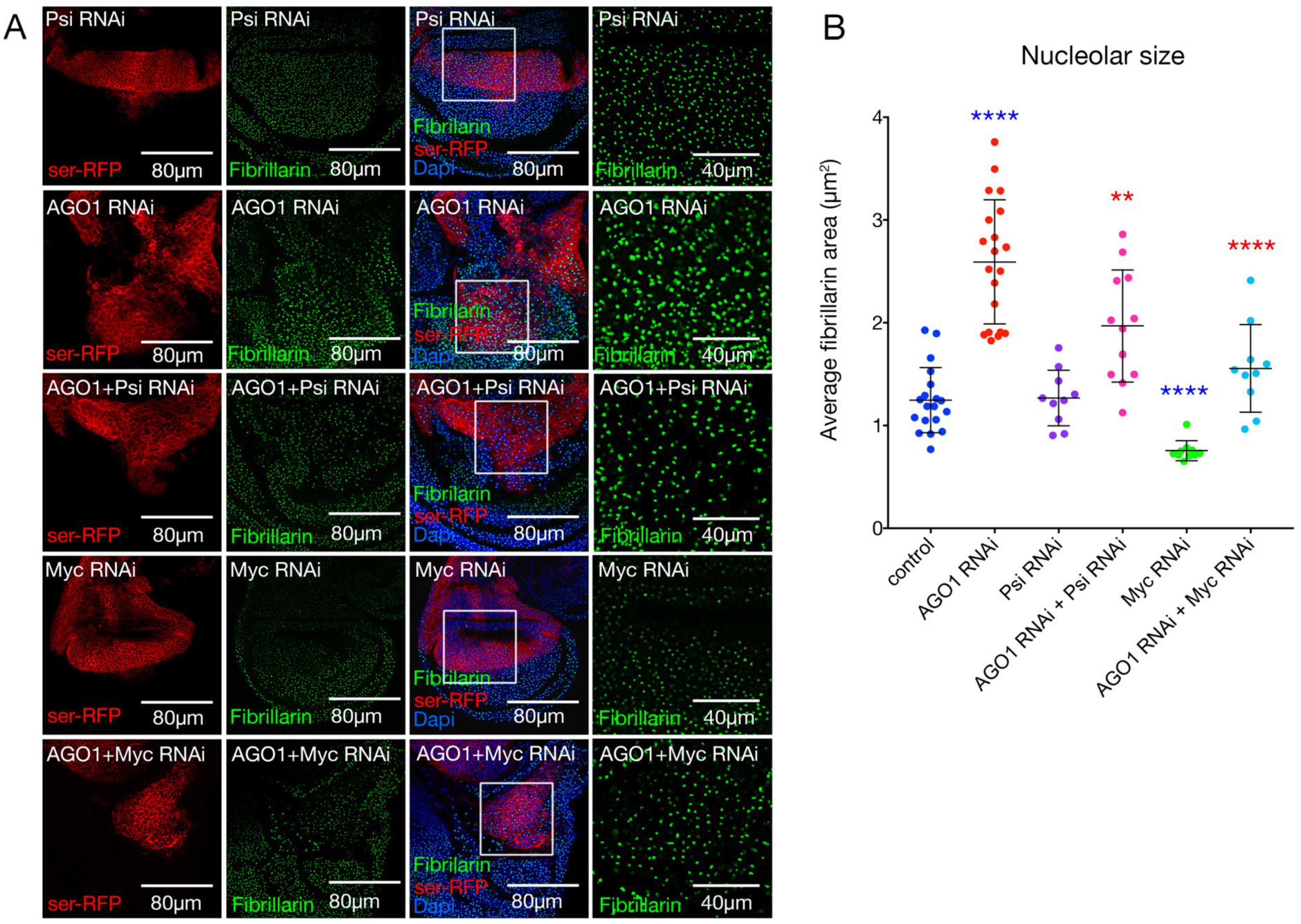
Increased nucleolar size due to AGO1 knockdown is dependent on Psi and Myc. **(A)** Control and *ser*-GAL4 driven RNAi in RFP-labelled cells for the genotypes marked in wing discs stained with anti-fibrillarin (green), DNA (blue). Zoom of region in the white square in the panel on the far right. **(B)** Quantification of average nucleolar area of approximately 50-100 nucleoli within the dorsal compartment for genotypes as marked, taken from a confocal z-section through the wing. Psi or Myc co-depletion significantly reduced nucleolar size in the *AGO1* RNAi background (p<0.0001 and p=0.0065 for Myc and Psi, respectively, compared with AGO1 knockdown alone).

### AGO1 depletion increases Myc abundance and function

The overgrowth observed in AGO1 knockdown wing imaginal disc cells (**Figure 3, 4**) was associated with a significant increase in *Myc* mRNA, which was reduced by Psi co-knockdown (**Figure 5A**). AGO1 knockdown also increased *Psi* mRNA (**Figure 5B**) and protein abundance (**Supplementary Figure 3**), suggesting AGO1 might increase Myc (at least in part) by increasing abundance of Psi. Together, these data suggest AGO1 represses *Myc* expression in larval wing discs in a manner partially dependent on Psi. In accordance with AGO1 normally being required for *Myc* repression, AGO1 depletion also increased Myc protein levels (**Figure 5C**). To determine whether increased *Myc* mRNA and protein resulted in heightened Myc function (i.e. transcriptional activity) we monitored abundance of two established Myc target genes in mammalian and *Drosophila* systems, *Polr1c* (polymerase I polypeptide C) and *Cad* (carbamoyl-phosphate synthetase 2) (Mitchell et al., 2015; Poortinga et al., 2014; Poortinga et al., 2011). *Polr1c* and *Cad* mRNA were significantly increased following AGO1 depletion and co-knockdown of Psi or Myc reduced abundance of these Myc target mRNAs (**Figure 5D-E**). Together, these data suggest AGO1 is essential for restraining *Myc* levels and preventing cell overgrowth.

**Figure 5.**
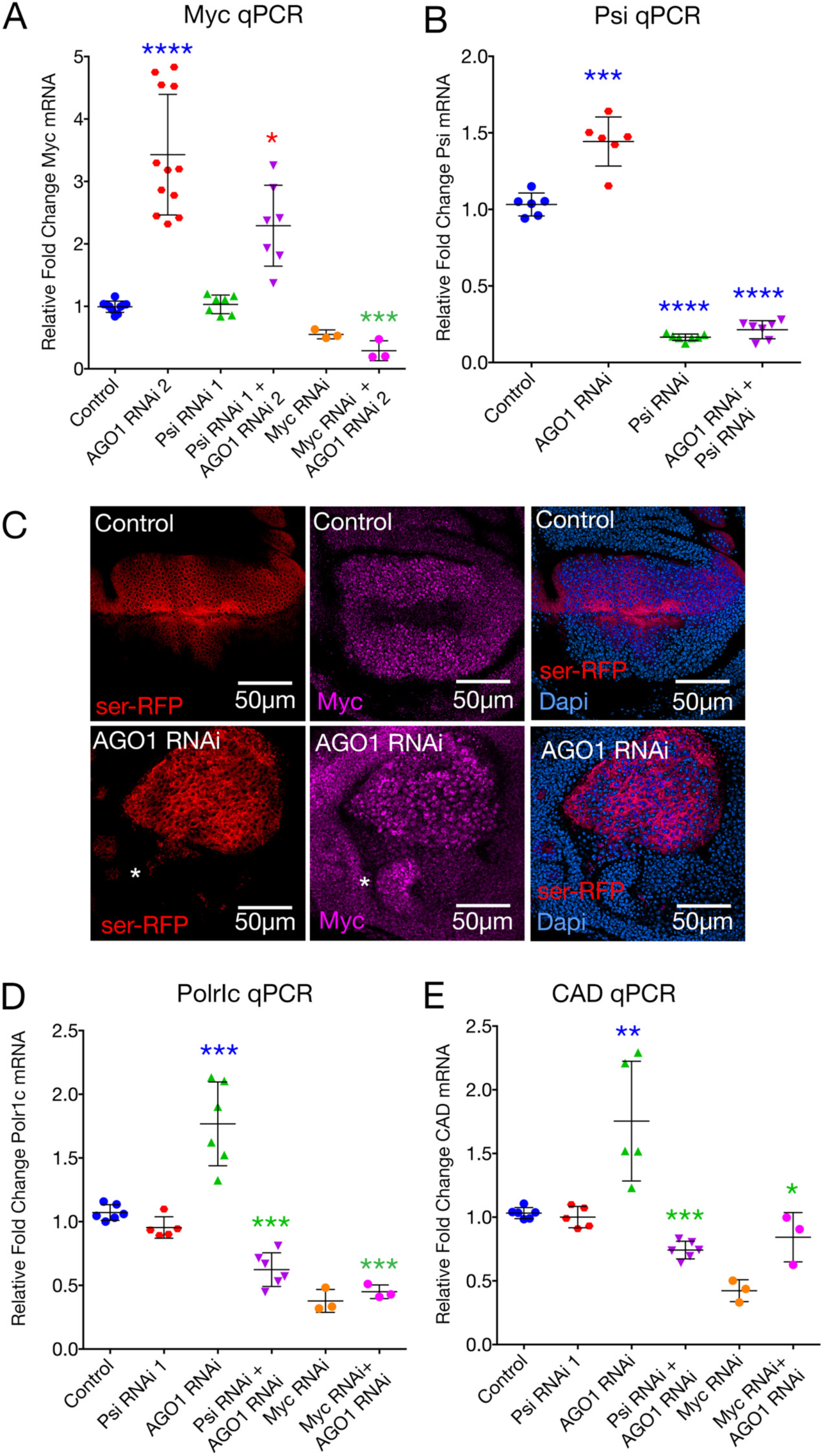
AGO1 knockdown increases Myc abundance and function in larval wing discs. **(A)** *Myc* qPCR following *AGO1* RNAi and/or *Psi* RNAi knockdown. AGO1 knockdown significantly increased *Myc* mRNA, compared with control (p<0.0001). Psi co-knockdown significantly reduced *Myc* mRNA compared with *AGO1* RNAi alone (p=0.0134). **(B)** *Psi* qPCR for *AGO1* RNAi and/or *Psi* RNAi knockdown wing discs. AGO1 knockdown increased *Psi* mRNA compared with control (p=0.0002). **(C)** Anti-Myc antibody (purple) on *ser-GAL4* driven *AGO1* RNAi compared with control. A region with elevated Myc protein in non-AGO1 knockdown cells marked with (*) **(D, E)** qPCR for Myc-target genes (*Polr1c* and *CAD*) in wing discs following *AGO1* RNAi and/or *Psi* RNAi knockdown. AGO1 knockdown significantly increased *Polr1c* and *CAD* mRNA, compared with control (p=0.0005 and p=0.043, respectively). Psi co-knockdown significantly reduced *Pol1rc* and *CAD* mRNA compared with *AGO1* RNAi alone (p<0.0001 and p=0.0005, respectively).

### Neither miR-996 nor miR-308 drive *Myc* mRNA turnover

As AGO1 induces miRNA-dependent mRNA degradation as part of the RISC complex (Hutvagner and Simard, 2008), we tested whether AGO1 depletion altered *Myc* mRNA levels post transcriptionally. We screened miRBase (Griffiths-Jones, 2004), which contains published mature miRNA sequences from 223 species (Kozomara and Griffiths-Jones, 2014), for miRNAs predicted to target the *Myc* 3’UTR by sequence similarity (http://www.mirbase.org). miR-308 and miR-996 were the only miRNAs predicted to target *Myc* (**Supplementary Figure 4A**) that were also expressed in third instar larval tissues based on the modENCODE database (Contrino et al., 2012). In *Drosophila* embryos miR-308 drives *Myc* mRNA and protein depletion (Daneshvar et al., 2013), however overexpression of miR-308 did not reduce *Myc* mRNA in the larval wing imaginal disc (**Supplementary Figure 4B)**, suggesting mir-308’s capacity to target *Myc* depends on developmental context. In contrast, miR-996 overexpression significantly increased *Myc* mRNA abundance (**Supplementary Figure 4B)**, indicating *Myc* mRNA is not a target for miR-996 driven degradation in the wing. Moreover, the capacity of *AGO1* knockdown to increase *Myc* was not altered by miR-308 nor miR-996 overexpression (**Supplementary Figure 4B)**, suggesting AGO1 repression of *Myc* is not dependent on the function of either of the miRNAs predicted to target *Myc.*

### AGO1 protein localises to the cytoplasm and the nucleus

AGO proteins, together with some components of the RISC, have been reported to enter the nucleus and regulate transcription (Catalanotto et al., 2016; Gosline et al., 2016; Kalantari et al., 2016; Shimada et al., 2016; Thomson et al., 2015; Woolnough et al., 2015). In early stage *Drosophila* blastoderm embryos, AGO1 protein localises to both the nucleus and cytoplasm (Pushpavalli et al., 2012). Biochemical fractionation and confocal immunofluorescence have also detected AGO proteins in the nuclear compartment of mammalian cells (Ahlenstiel et al., 2012). We therefore investigated the localisation of AGO1 in wing imaginal disc cells using an anti-AGO1 antibody and an AGO1-GFP protein trap, which generates a GFP fusion with endogenous AGO1 (Buszczak et al., 2007). As expected, given miRNA silencing functions, AGO1 and the AGO1-GFP fusion localised predominantly to the cytoplasm in both the wing discs (**Figure 6A and B**) and salivary glands (**Supplementary Figure 5**). In addition, co-staining with lamin to mark the nuclear envelope revealed punctate AGO1 staining within the nucleus.

**Figure 6.**
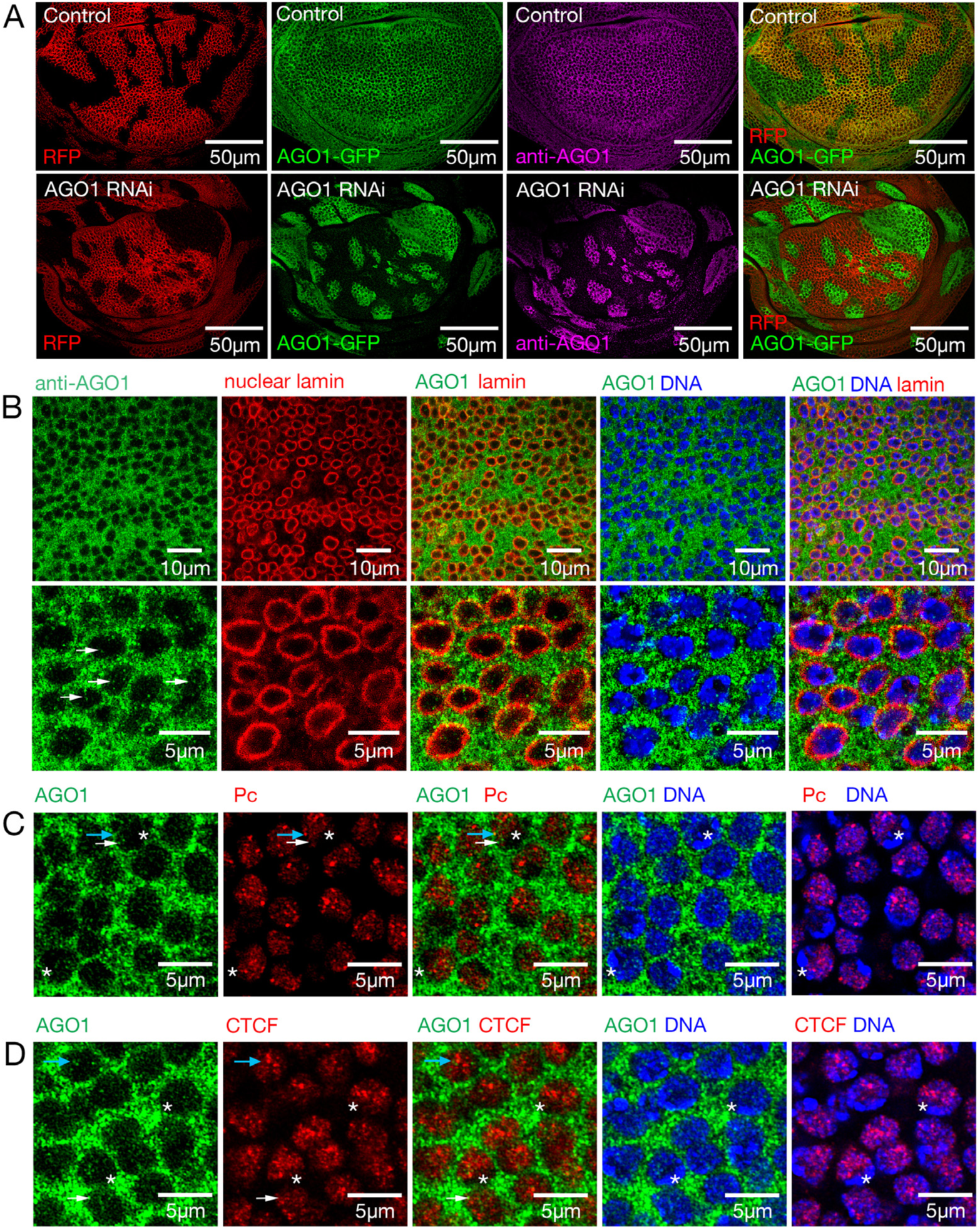
AGO1 protein localises to both cytoplasmic and nuclear compartments. **(A)** RFP-marked control and *AGO1* RNAi flip out clones in wing imaginal discs 3 days after heatshock in the AGO1-GFP protein trap background, stained with AGO1 antibody (purple). Wild type third instar wing imaginal discs stained with **(B)** AGO1 antibody (green) and nuclear lamin (red) (**C**) Single 1 µm Z section from Zeiss Airyscan images for AGO1 antibody (green) and Pc-GFP (false coloured red). Blue arrow marks AGO1 puncta with PcG overlap, white arrow marks AGO1 without clear PcG overlap. Asterisks mark heterochromatin with intense Dapi staining. (**D**) AGO1 (green) and CTCF-GFP (false coloured red) for single 1 µm Z sections. A blue arrow marks 3 AGO1 puncta, the closest 2 without and furthest one with CTCF overlap. White arrow marks AGO1 with weak CTCF overlap. Asterisks mark heterochromatin with intense Dapi staining.

As previous analysis in *Drosophila* antennal discs reported substantial overlap between AGO1 protein and Polycomb body foci (42% colocalization) (Grimaud et al., 2006), we examined whether AGO1 localises to regions of PcG silencing in wing imaginal discs by staining with anti-AGO1 in the Pc-GFP background to mark PcG foci (Dietzel et al., 1999). In contrast to the earlier studies using lower resolution microscopy, our high-resolution analysis separated PcG bodies from AGO1 puncta, revealing limited direct overlap (**Figure 6C**). Indeed, quantification revealed overlap of just 8% and close proximity of 8.4% between AGO1 and PcG complexes, with the majority (83.6%) of staining occurring independently (**Supplementary Figure 6**). To confirm that PcG bodies overlap euchromatin, as previously reported (Pirrotta and Li, 2012), Dapi was used to distinguish heterochromatin by intense staining, which revealed both PcG bodies and AGO1 puncta in regions of weaker Dapi staining i.e. AGO1 and PcG localise with euchromatin (**Figure 6C**).

The observation that AGO1 puncta and PcG bodies localise to euchromatin, but generally do not directly overlap (83.6%), lends support to the idea that multi-protein and RNA complexes comprising AGO1 might serve as a scaffold for assembly of the PcG super-complexes, that underlie both PcG and insulator bodies (Pirrotta and Li, 2012; Shevtsov and Dundr, 2011). Although AGO2 has been reported to enable insulator function independent of RNAi activity through physical association with CTCF binding sites in *Drosophila* (Moshkovich et al., 2011), such roles have not been reported for AGO1. We therefore tested proximity between AGO1 and chromatin insulator bodies, and the localisation of AGO1 and CTCF chromatin insulator complexes in the nucleus using anti-AGO1 and CTCF-GFP (**Figure 6D**). As expected, based on Dapi staining, AGO1 complexes were detected in regions of euchromatin, however, only 15% of the AGO1 puncta were found overlapping or in close proximity with CTCF-marked insulator domains (**Figure 6D**, quantified **Supplementary Figure 6**). Together these data suggest AGO1 complexes interact with a small subpopulation of PcG transcriptional silencing loci and CTCF insulator domains in the nucleus.

### AGO1 knockdown increases *Myc* transcription

Recent studies demonstrated transcriptional regulation of the *MYC* oncogene involving looping of super enhancers and the *MYC* promoter requires a conserved CTCF site (Schuijers et al., 2018). This, together our observations that AGO1 interacts with the RNA Pol II machinery, localises to euchromatic regions of DNA and overlaps PcG and CTCF complexes, led us to investigate whether AGO1 regulates *Myc* mRNA abundance at the level of transcription. Indeed, AGO1 is required to constrain the *Myc* promoter, as *Myc-lacZ* enhancer trap (Mitchell et al., 2010; Peter et al., 2002) activity was significantly increased in the *AGO1* knockdown wing disc compartment (**Figure 7A**). To further investigate whether increased *Myc* mRNA associated with AGO1 loss of function was due to altered transcription, we designed primers to the first intron of *Myc* to measure pre-mRNA levels. qPCR revealed an increase in the immature *Myc* message following AGO1 knockdown in wing discs (**Figure 7B**). Together these data suggest the increased *Myc* expression associated with AGO1 depletion is due to activation of the *Myc* promoter and increased transcription.

**Figure 7.**
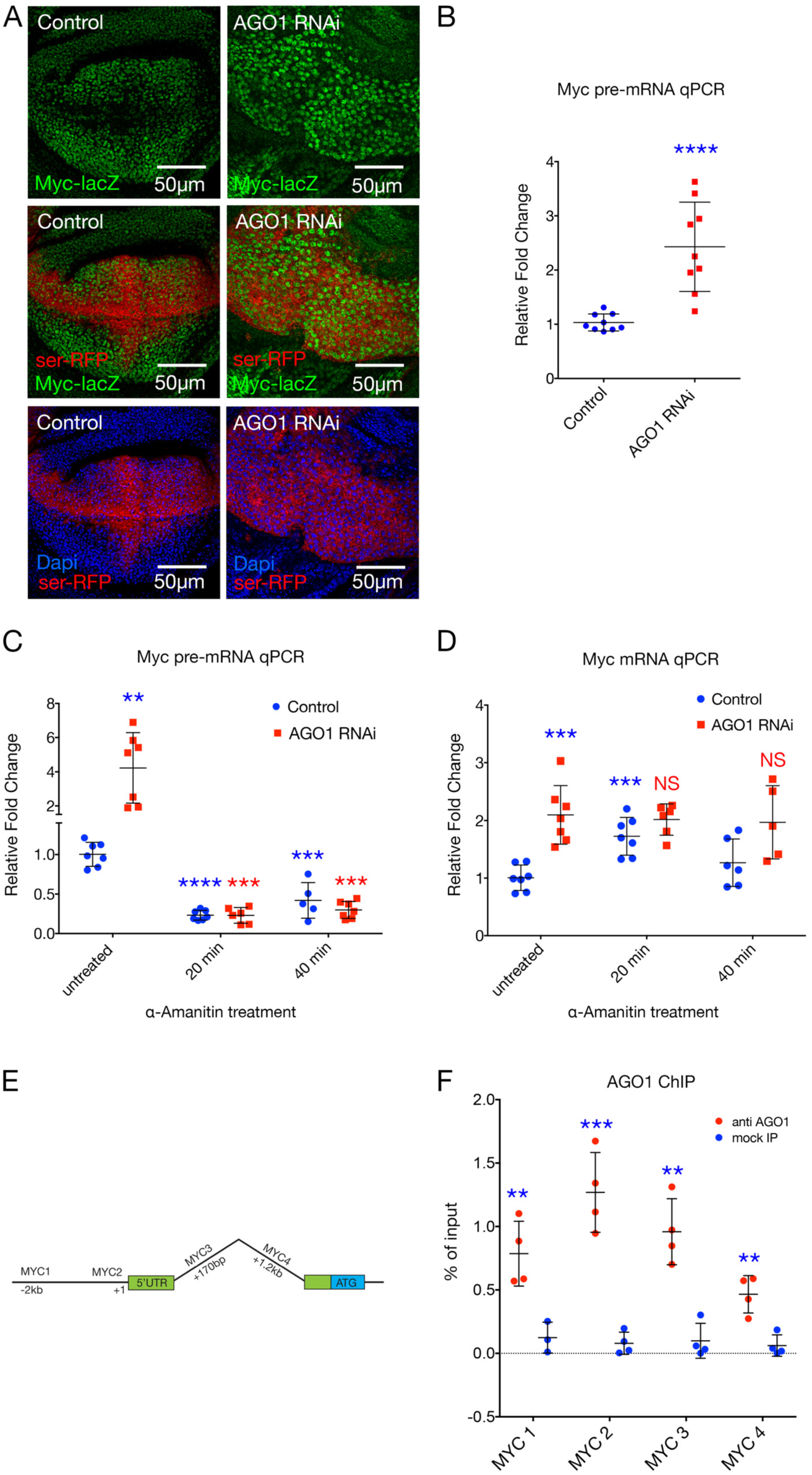
AGO1 represses *Myc* at the level of transcription. **(A)** *Myc-lacZ* enhancer trap activity, marked with anti-β-Gal antibody (green) for *ser*-GAL4 driven *AGO1* RNAi in the *UAS-*p35 background compared with *UAS-*p35 alone control, marked with RFP and stained for DNA (blue). **(B)** qPCR for *Myc* pre-mRNA following *AGO1* RNAi knockdown in larval wing discs, p<0.0001 compared with control. **(C)** qPCR for *Myc* pre-mRNA following AGO1 knockdown compared with control for larval head tissues treated with α-amanitin for 0 min, 20 min and 40 min or untreated, as marked. p<0.001 for untreated compared with α-amanitin treated *AGO1* RNAi at either time point **(D)** qPCR for mature *Myc* mRNA, genotypes and treatment with α-amanitin as marked. **(E)** Schematic of *Myc* showing the position of the amplicons used for qPCR. **(F)** AGO1 ChIP on wild type larval tissues compared to no-antibody control (p=0.0003 for AGO1 antibody compared with input).

### Increased *Myc* due to AGO1 depletion requires RNA Pol II transcription

To determine if AGO1 regulates *Myc* expression at the transcriptional level we used α-Amanitin to block RNA Pol II activity. Consistent with observations using wing imaginal discs (**Figure 5**), *Myc* pre- and processed mRNA levels were significantly increased in untreated larval head tissues following AGO1 depletion (**Figure 7C-D**). Interestingly, although *Myc* pre-mRNA was significantly decreased 20 minutes and 40 minutes after α-Amanitin treatment (**Figure 7C**), mature mRNA was significantly increased in control tissues at the 20-minute time point (**Figure 7D**), suggesting *Myc* mRNA stability might increase in response to transcriptional inhibition. In the AGO1 knockdown background, *Myc* pre-mRNA levels were dramatically decreased following α-Amanitin treatment (**Figure 7C**). Thus, the AGO1 knockdown-induced increase in *Myc* pre-mRNA was dependent on RNA Pol II transcriptional activity. Abundance of the mature *Myc* transcript was not significantly altered in the AGO1 knockdown background following α-Amanitin treatment, again suggesting feedback mechanisms might result in increased mRNA stability in response to RNA Pol II inhibition. Together with the observation that AGO1 knockdown increased *Myc* promoter activity in the wing discs, these data suggest AGO1 is normally required for repression of *Myc* transcription.

### AGO1 is enriched on *Myc*

The increased *Myc* promoter activity following AGO1 depletion, and increased *Myc* mRNA abundance requiring RNA Pol II activity led us to test whether AGO1 directly regulates *Myc* transcription. We therefore performed ChIP using the anti-AGO1 antibody followed by qPCR with amplicons flanking the *Myc* transcription start site (**Figure 7E**). In wild type larval tissues, significant AGO1 enrichment was observed in *Myc* regulatory regions compared with the mock-IP control, with highest enrichment observed at the transcription start site (**Figure 7F**), suggesting AGO1 normally inhibits *Myc* transcription through direct interaction with the *Myc* promoter.

## Discussion

Here we demonstrate a novel role for AGO1 as a growth inhibitor in *Drosophila.* AGO1 depletion was sufficient to increase Myc (mRNA and protein) to drive ribosome biogenesis, nucleolar expansion, and cell growth in a Myc and Psi-dependent manner. The increased *Myc* promoter activity in AGO1 knockdown wing discs, together with the *α*-Amanitin-dependent increase in *Myc* pre-mRNA abundance, suggests AGO1 represses *Myc* at the level of transcription. In accordance with the AGO1’s observed growth inhibitory capacity, in AGO1 knockdown wings increased *Myc* mRNA and protein abundance were associated with increased Myc function i.e. activation of established Myc targets. Interestingly, although Psi knockdown only modestly decreased *Myc* mRNA levels in AGO1-depleted wings, Psi co-depletion strongly reduced expression of Myc targets. This observation suggests Psi is not only required for *Myc* transcription (Guo et al., 2016) but may also be required for activation of Myc growth targets in the context of AGO1 depletion. Thus, future studies are required to determine whether Psi and Myc bind common targets and if Psi is required for transcriptional activation of Myc target genes.

Recent genome-wide functional RNAi screens in *Drosophila* S2 cells, identifying AGO1 as a modifier of Polycomb foci, suggested extra-miRNA functions for AGO1 (Gonzalez et al., 2014). PcG mediates epigenetic repression of key developmental genes to control cell fate, and PcG repression is stabilised via aggregation of PcG foci in the nucleus. AGO1 depletion disrupted nuclear organization and reduced intensity of Pc foci, suggesting AGO1 negatively regulates PcG-mediated silencing (Grimaud et al., 2006). The *Drosophila* PcG complex has been characterised for roles in silencing homeotic genes by binding Polycomb group response elements (PREs), including the Fab-7 PRE-containing regulatory element from the Hox gene, Abdominal-B. Components of the RNAi machinery, including AGO1 and Dicer-2, have been implicated in driving PcG-dependent silencing between remote copies of the Fab-7 element, engineered throughout the genome to monitor long-distance gene contacts. Interactions between Hox genes silenced by PcG proteins were decreased in *AGO1* mutants, suggesting AGO1 regulates nuclear organisation, at least in part, by stabilising PcG protein recruitment to chromatin (Grimaud et al., 2006).

The question remains regarding how AGO1 targets *Myc* transcription. The physical and genetic interaction between Psi and AGO1, and observation that AGO1 loss-of-function mutants restore cell and tissue growth in the Psi knockdown wing, suggests AGO1 inhibits growth dependent on this Myc transcriptional regulator. AGO2 has been implicated in insulator-dependent looping interactions defining 3D transcriptional domains (TADs) through association with dCTCF binding sites in *Drosophila* (Moshkovich et al., 2011). Although similar roles for AGO1 have not been reported, the cancer-related super-enhancers for the *MYC* oncogene lie within the 2.8 Mb TAD and control *MYC* transcription via a common and conserved CTCF binding site located 2 kb upstream of the *MYC* promoter i.e. in proximity with the FUSE (1.7kb upstream) bound by FUBP1. Moreover, gene disruption of the enhancer-docking site reduces CTCF binding and super-enhancer interaction, which results in reduced *MYC* expression and proliferative cell growth (Schuijers et al., 2018). AGO1-ChIP revealed significant enrichment on the *Myc* promoter, suggesting AGO1 likely interacts with Psi and the RNA Pol II machinery to directly regulate *Myc* transcription. Given the high level of conservation between AGO and CTCF proteins throughout evolution, it will be of great interest to determine whether human AGO1 also interacts with FUBP1 to regulate transcription of the *MYC* oncogene.

Here we have shown AGO1 behaves as a tumour suppressor during *Drosophila* development, through the ability to suppress *Myc* transcription, ribosome biogenesis and cell growth in the wing disc epithelium. Consistent with AGO1 having tumour suppressor activity, across a wide range of human cancers, large scale genomics data (cBioPortal (Gao et al., 2013)) identified AGO1 as frequently mutated or deleted in a diverse variety of tumours (e.g. reproductive, breast, intestinal, bladder, and skin cancers). Region 1p34–35 of chromosome 1, which includes AGO1, is frequently deleted in Wilms’ tumours and neuro-ectodermal tumours (Parisi et al., 2011). In neuroblastoma cell lines, AGO1 behaves as a tumour suppressor, with overexpression heightening checkpoint sensitivity and reducing cell cycle progression, and GEO Profile microarray data inversely correlates AGO1 expression with proliferative index (Parisi et al., 2011) i.e. AGO1 levels are significantly lower in tumorigenic cells compared with differentiated cells (Barrett et al., 2011). In the context of cancer, it will be important to determine whether AGO1 loss-of-function alters *MYC-*dependent cancer progression and vice versa. As increased abundance of the MYC oncoprotein is associated with the pathogenesis of most human tumours (Dang, 2012; Levens, 2010), deciphering such novel mechanisms of *MYC* repression will be fundamental to understanding MYC-dependent cancer initiation and progression.

## MATERIALS AND METHODS

### *Drosophila* Strains

The *UAS-AGO1* RNAi 1 (BL53293), *UAS-AGO1* RNAi 2 (BL33727), *UAS-Myc* (BL9675), AGO-GFP (BL50805), *UAS-miR-308* (BL41809), *UAS*-*miR-996* (BL60653), *Myc-lacZ* (BL12247), CTCF-GFP (BL64810), Pc-GFP (BL9593) *ser-*GAL4 (BL6791), *tub*-GAL4 (BL5138) and *tub*-GAL80ts (BL7019) were obtained from the Bloomington Stock Centre. The *UAS-Myc* RNAi line (V2947), *UAS-Psi* RNAi 1 (V105135), *UAS-Psi* RNAi 2 (V28990), *AGO1*^*k00208*^ (V10470) and *AGO1*^*04845*^ (V11388) lines were obtained from the Vienna *Drosophila* RNAi Center.

### Co-Immunoprecipitation and Western Blotting

Co-Immunoprecipitation (Co-IP) was performed using 25 wild type 3rd instar larval heads dissociated in cold lysis buffer (50 mM Tris pH 7.5, 1.5 mM MgCl2, 125 mM NaCl, 0.2% NP40, 5% glycerol, 1x Protease inhibitor cocktail). Following homogenization, protein was collected by centrifugation at 12 000 rpm for 10 min at 4°C. The extract was pre-cleared by incubation with nProtein A Sepharose ™ beads (GE Healthcare Life Science) for 1 hour at 4°C with rotation and the supernatant collected by centrifugation at 12 000 rpm. Equal amounts of pre-cleared protein lysate were incubated with either anti-AGO1 (Abcam, ab5070) or anti-Psi (custom generated rabbit polyclonal antibody, Biomatik) antibodies overnight at 4°C. Beads were washed with lysis buffer 5 times, and the eluent resolved using 10% SDS PAGE/Western with appropriate primary antibody prior to detection with Li-Cor Odyssey IR detection.

### Immunofluorescence, microscopy and image analysis

Wandering 3^rd^ instar larvae were dissected and fixed for 20 min in 4% paraformaldehyde (PFA), washed in PBS with 0.1% Tween (PBT), blocked in 5mg/ml Bovine Serum Albumin (BSA) prior to incubation overnight at 4°C with primary antibody. Due to the high level of cell death and lethality associated with constitutive *ser*-GAL4 driven AGO1 depletion, analysis with AGO1 antibody was conducted in the temperature sensitive *tubulin*-GAL80 (*tub*-GAL80ts) background 24 hours after GAL4 induction. Primary antibodies used for immunofluorescence; Myc N (Rabbit 1 in 500 Santa Cruz d46-507), Fibrillarin (Rabbit 1 in 500 Abcam ab5821), AGO1 (Rabbit, 1 in 500 Abcam ab5070), Psi (Rabbit 1 in 500 custom-made via Biomatik) Lamin (Mouse, 1 in 20, DSHB ADL 101), β-galactosidase (Chicken 1 in 1000, Abcam ab9361). After incubating with appropriate fluorophore-tagged secondary antibodies samples were counterstained with DAPI solution and wing imaginal discs imaged using the Zeiss LSM800 confocal microscope (Zen Blue software). Overlapping 1 um Z-sections were collected at 40x magnification. Fluorophores were imaged using band-pass filters to remove cross-detection between channels. Images were processed and prepared using Image J and Adobe Photoshop CS5. Fibrillarin was quantified in FIJI on confocal z-sections of wing columnar epithelial cells, merged to display maximum projections (2-3 sections). Thresholding was performed and images were used to measure average Fibrillarin area in the dorsal compartment marked by *serrate-*GAL4>*UAS*-RFP expression. 50-100 nucleoli were selected using freeform selection tool, and analysed with the “Analyse Particles” tool, with minimum particle size of 0.5 µm^2^ applied in order to exclude noise and out of focus nucleoli. The output used image metadata to calculate average nucleolar area in µm^2^ for each wing disc analysed. % Overlap between AGO1 and PcG/CTCF were performed in FIJI by thresholding to isolate individual puncta and overlaying channels to detect co-occurrence or adjacency, which was counted and expressed as proportion of total AGO1 puncta per individual nuclei.

### Adult wing size analysis

Adult wing size was determined for male wings imaged with an Olympus SZ51 binocular microscope, at 4.5x magnification using the Olympus DP20 camera. Wing size was measured by pixel count for the area posterior to wing vein L5, using Photoshop software CS5. For wing hair counts, adult male wings were imaged with Olympus BX 61 microscope at 20x magnification using the Olympus DP70 camera. Wing cell size was determined using wing hair counts in a defined area (200 × 100 pixels) at the central region posterior of wing vein L5. Then the hair number was converted to relative single hair/cell size via dividing the area of the fixed region by hair numbers.

### qPCR

RNA was isolated from equivalent numbers of wing imaginal discs (10 pairs for each genotype) using the Promega ReliaPrep RNA Cell miniprep system and eluted in 20 uL nuclease-free water. RNA purity and integrity were assessed using an automated electrophoresis system (2200 TapeStation, Agilent Technologies). 6 uL of RNA was used for each cDNA synthesis (GoScript™ Reverse Transcription System kit, Promega). qPCR was performed using Fast SYBR Green Master Mix (Applied Biosystems) using the StepOnePlus Real-Time PCR System and Sequence Detection Systems in 96-well plates (Applied Biosystems, 95°C for 2 min, 40 cycles 95°C 1 s and 60°C 20 s). Amplicon specificity was verified by melt curve analysis. Average Ct values for two technical replicates were calculated for each sample. Multiple internal control genes were analyzed for stability and target gene expression was normalized to the mean of *cyp1* and *tubulin* or *cyp1* alone, selected for having high expression and little sample-to-sample variability as determined by RefFinder. Fold change was determined using the 2-ΔΔ CT method.

### Primers used for qPCR

Myc

5’ GTGGACGATGGTCCCAATTT 3’

5’ GGGATTTGTGGGTAGCTTCTT 3’

Psi

5’ CGATGGCATCCCATTTGTTTGT 3’

5’ GGTGGTCAAGACTACTCGGC 3’ AGO1

5’ ACTCTACGGTCTGTCCGTTC 3’

5’ CCCGCTCAGATGCAATCATTC 3’

5’ETS

5’ GGCAGTGGTTGCCGACCTCG 3’

5’ GCGGAGCCAAGTCCCGTGTT 3’

Tubulin

5’ TGGGCCCGTCTGGACCACAA 3’

5’ TCGCCGTCACCGGAGTCCAT 3’

CYP1

5’ TCGGCAGCGGCATTTCAGAT 3’

5’ TGCACGCTGACGAAGCTAGG 3’

Pol r1c

5’ TGTATCCCGCCATTGCAA 3’

5’ GGGCACATCGCTGAGCAF 3’

Cad

5’ CATTGGCAGTTTCAAGCACAA 3’

5’ TCTTGGCCAGATCCCGTATG 3’

### α-Amanitin treatment

α-Amanitin inhibits RNA Pol II-dependent transcription, therefore interfering with mRNA production (Lindell et al., 1970). α-Amanitin (sigma #A2263) was diluted in 1 mL of Nano pure water to make a 1mg/mL stock solution, which was stored at −20°C in the dark. Third instar larval heads were dissected and incubated with freshly made 20 ug/mL α-Amanitin in Schneider’s Medium at 25°C for 0 min, 20 min, 40 min respectively. After α-Amanitin treatment, samples were washed for 5 minutes using fresh Schneider’s Medium and snap frozen in 250 uL LBA+TG lysis buffer from the Promega ReliaPrep RNA Cell miniprep kit. Following RNA extraction and cDNA synthesis, qPCR was performed and analysed (as above 2.3.1) with *Myc* cDNA primers and *Myc* pre-mRNA primers.

### Primers for *Myc* pre-mRNA qPCR

Myc pre-mRNA

5’ TTCAAAATAGAATTTCTGGGAAAGGT 3’

5’ GCGGCCATGATCACTGATT 3’

### Chromatin Immunoprecipitation

Chromatin immunoprecipitation (ChIP) assays were carried out as described previously (Lee et al., 2015). Briefly, for each ChIP sample 30 larval heads were collected from mid 3^rd^ instar larvae and fixed in 4% PFA for 20 minutes. Larval heads were dissociated and chromatin sheared in 0.4% sodium dodecyl sulphate (SDS) using a Covaris S2 (10 min duration, 10% DUTY, 200 cycles per burst, Intensity 4, achieving average DNA fragment sizes 200–600 bp). ChIP was performed in IP buffer containing 0.1% SDS and 3 μg of antibody was used for each IP (anti-RNA Pol II phospho S5 antibody (ab5131), or anti-RNA Pol II phosphor S2 (ab5095). Analysis was performed in triplicate using Fast SYBR Green Master Mix (Applied Biosystems) on a StepOnePlus Real-Time PCR System and Sequence Detection Systems in 384-well plates (Applied Biosystems). To calculate the percentage of total DNA bound, non-immuno precipitated input samples from each condition were used as the qPCR reference for all qPCR reactions.

### Primers for ChIP qPCR

MYC 1

5’ GGCGATCGTTTCTGGCCTACGG 3’

5’ GCAGGCGCATTTGACTCGGC 3’

MYC 2

5’ ACTACTACTAACAACTGTCACAAGCCAAGT 3’

5’ TTTATGTATTTGCGCGGTTTTAAG 3’

MYC 3

5’ TTCAAAATAGAATTTCTGGGAAAGGT 3’

5’ GCGGCCATGATCACTGATT 3’

MYC 4

5’ GGTTTTCCTTTTATGCCCTTG 3’

5’ CTATTAACCATTTGAACCCGAAATC 3’

### Statistical analysis

All statistical tests were performed with Graphpad Prism 6 using unpaired 2-tailed t-test with 95% confidence interval. In all figures error bars represent SD and according to the Graphpad classification of significance points * (P = 0.01–0.05), ** (P = 0.001–0.01), *** (P = 0.0001– 0.001) and **** (P< 0.0001).

